# Developing a More Accurate Biomedical Literature Retrieval Method using Deep Learning and Citations in PubMed Central Full-text Articles

**DOI:** 10.1101/2021.10.21.465340

**Authors:** Chun-chao Lo, Shubo Tian, Yuchuan Tao, Jie Hao, Jinfeng Zhang

## Abstract

Most queries submitted to a literature search engine can be more precisely written as sentences to give the search engine more specific information. Sentence queries should be more effective, in principle, than short queries with small numbers of keywords. Querying with full sentences is also a key step in question-answering and citation recommendation systems. Despite the considerable progress in natural language processing (NLP) in recent years, using sentence queries on current search engines does not yield satisfactory results. In this study, we developed a deep learning-based method for sentence queries, called DeepSenSe, using citation data available in full-text articles obtained from PubMed Central (PMC). A large amount of labeled data was generated from millions of matched citing sentences and cited articles, making it possible to train quality predictive models using modern deep learning techniques. A two-stage approach was designed: in the first stage we used a modified BM25 algorithm to obtain the top 1000 relevant articles; the second stage involved re-ranking the relevant articles using DeepSenSe. We tested our method using a large number of sentences extracted from real scientific articles in PMC. Our method performed substantially better than PubMed and Google Scholar for sentence queries.

## Introduction

Literature retrieval using search engines is routinely performed by biomedical scientists to find academic papers (related to a set of keywords) they are interested in. Public search engines, such as PubMed and Google Scholar, have been commonly used for this purpose. The ever-increasing number of biomedical research articles published every year has made it very challenging for search engines to rank the most relevant articles highly enough for users to find them. Missing important studies in a literature search can have serious consequences when designing a new study, such as wasting resources and/or time; it can also result in making wrong conclusions or missing new discoveries when interpreting experimental results.

The importance of search engines for scientific literature retrieval has motivated researchers to develop effective solutions over the years, including both traditional methods [1-7] and more recently, deep learning methods [8-17]. Web-based tools have also been developed so that researchers can conduct a literature search from various databases [18]. Most search engines use BM25 (or a variation of it) for its similarity ranking algorithm [19]. BM25 is a bag-of-words retrieval function that ranks a set of documents based on the number of query terms appearing in each document, regardless of their proximity within the document. A number of variations [20] have been proposed to address the known issues of BM25 [21]. User activity information has also been leveraged to improve users’ search experience [22-24]. In 2017, a new relevance search algorithm called “Best Match” was deployed at PubMed [24] using a “learn to rank” based machine learning algorithm [25-27] trained by user-click information from PubMed search logs. PubMed has been the major literature search tool for biomedical scientists around the world with more than 3 million visits per day [28, 29].

When searching for papers related to a particular topic of interest, it is more effective to search with more keywords that define a topic, such as using full sentences or asking questions. However, current search engines do not perform well for such query types (see Results for more details). For example, PubMed often does not return any results for sentence queries. In this study, we aim to develop a better method for document retrieval using sentence queries by taking advantage of recent advancements in deep learning algorithms for natural language processing (NLP). Methods that use sentence queries for document retrieval will also help the development of more accurate question-answering and citation recommendation systems.

A major challenge when developing a quality search engine using machine learning methods is the availability of a large amount of labeled training data. The labeled data should consist of different queries matched to relevant papers. Generating such labeled data manually is very time and resource consuming. To tackle this challenge, we used the citation data from PubMed Central (PMC) full-text articles to generate labeled data. In full-text articles, when a sentence cited an article, we call the sentence as a citing sentence and the article as a cited article. They can be considered as a manually labeled case with the citing sentence as the query and the cited article as the relevant document. We can extract citing sentences and the corresponding cited articles from PMC full-text articles to generate millions of labeled cases, which can be used to train quality deep learning models.

We tried several deep learning architectures and found that the decomposable attention model [30] offered the best tradeoff between accuracy and speed. We used a two-step approach, where in the first step a modified BM25 method was used to rank all the articles to generate the top 1000 relevant articles; then the articles were re-ranked in the second step using deep learning models to produce the final ranking.

We tested the performance of our method using a large number of cases obtained from the PMC citation data, which were not used for model training. Our method achieved significantly better performance than the search engines of PubMed and Google Scholar.

## Method and Data

### Problem formulation

Our goal is to find the most relevant articles associated with a sentence query from a database of articles. To use the latest deep learning methods, we need a substantial amount of labeled training data. To that end, we assume that the articles cited by a sentence in a scientific article are highly relevant to that sentence. Based on this assumption, we first downloaded full-text articles from PubMed Central (PMC) and MEDLINE citations from PubMed. We extracted sentences with citations from the full-text part of the PMC XML files, and obtained titles, abstracts, publication year, article type (journal article, review, case reports, etc.) and journal names from the MEDLINE citations to build an internal database. This internal database is necessary since we will need to query it many times to generate the training data. In addition, we downloaded journal citation reports from Web of Science [31] and extracted the impact factors as one of the input features. We also extracted citation information from PubMed XML files and calculated the number of citations an article has for as many articles as possible. For each citing sentence and the cited article associated with it, we created a sentence-article pair (SEN, ABS) as a true case, where ABS contains both the title and abstract of the cited article as well as additional information related to the cited article.

Applying a deep learning model directly to all the PubMed abstracts for a given query is not feasible due to the relatively high computational cost of deep learning models. To cope with this, we first built an SQL database of all the PubMed abstracts and used a modified BM25 algorithm [19] (called MBM) to query the top 1000 ranked articles, which are then re-ranked by the chosen deep learning model. When generating the training and validation dataset, we obtained two negative cases for each sentence query: one case was randomly chosen from the top 1,000 query results excluding the cited article, and the other case was randomly chosen from all the abstracts outside of the top 1000. This strategy allows the model to learn the general differences between the true and false cases and the subtle differences between the true and high-ranking false cases. When performing a sentence query, only articles published earlier than the article containing the citing sentence (SEN) were considered because SEN can cite only earlier articles. Our method is called DeepSenSe (Deep learning method using Sentence in Searches).

The workflow of our methodology development is shown in Figure 1.

**Figure 1.**
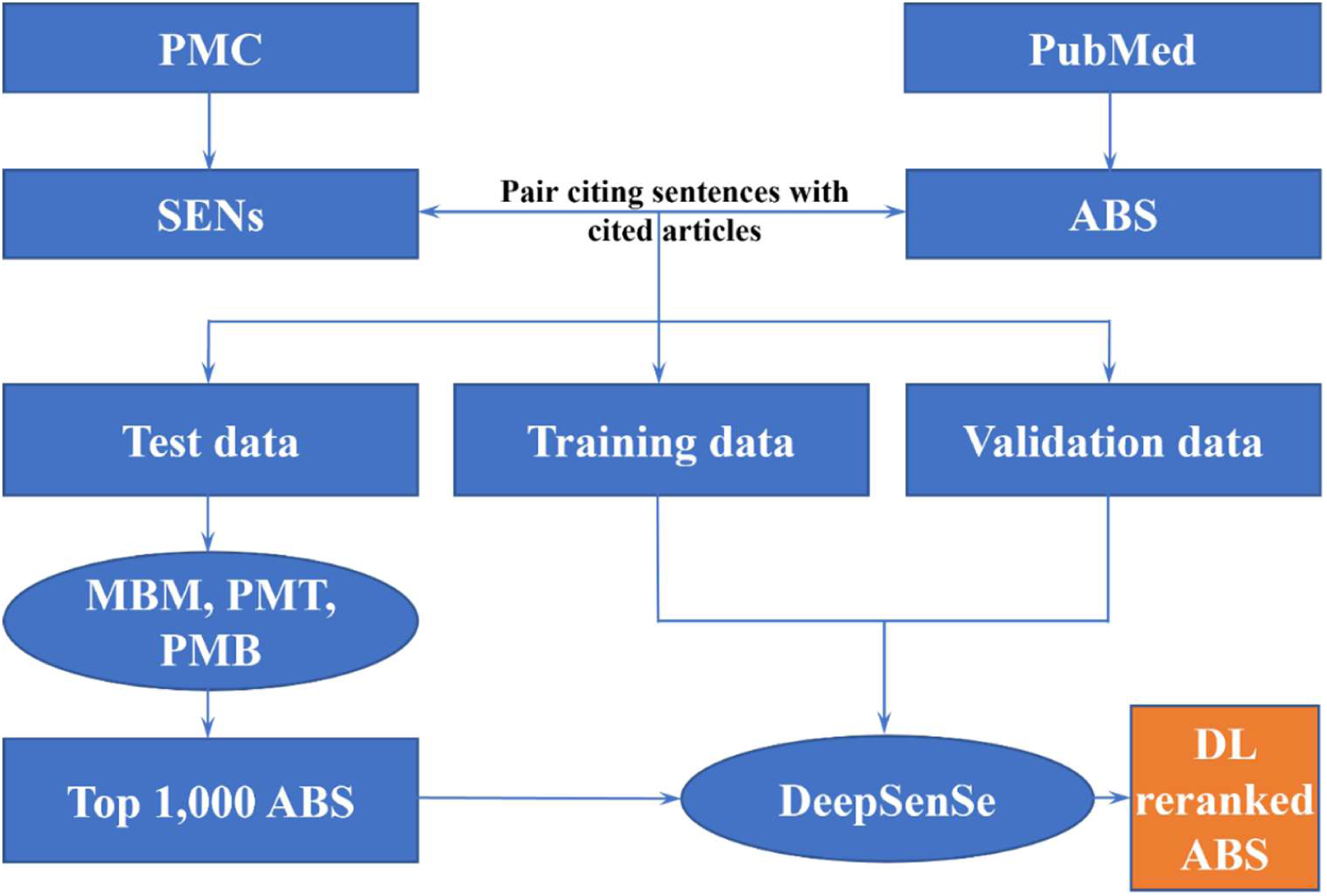
The workflow of our methodology development. SENs: citing sentences; ABS: titles and abstracts of cited articles. MBM: modified BM25 algorithm querying our database; PMT: PubMed TF-IDF algorithm querying PubMed database; PMB: PubMed “Best Match” algorithm querying PubMed database; DL: deep learning. Citing sentences were extracted from PMC full-text articles and split into training, validation, and test data. PubMed articles were used to build the MBM database. Citing sentences were paired with cited PubMed articles. In the training and validation data, the sentence-citation pairs served as true cases. False cases were created by randomly sampling two articles not cited by the sentence for each citing sentence. For testing, the citing sentence was used as the query sentence while the cited article was considered as the most relevant article. The citing sentences were used as queries to search using MBM, PMT, and PMB to get the top 1000 results which were then re-ranked by DeepSenSe to obtain the final ranking results.

### Datasets

We downloaded the PMC full-text articles published before Oct. 23rd, 2019. Sentences with citations were then extracted from the full-text. Sentences with no meaningful keywords were excluded. The remaining citing sentences were divided into training, validation, and test datasets. About 79% of citations are articles with a PubMed ID (PMID, a unique number for each article in PubMed); other citations are books, webpages, conferences, etc.

All the PubMed articles were stored and indexed in a MySQL database. A modified BM25 algorithm (MBM) [32] was used for querying articles from our own database [33]. The speed and accuracy of the MBM are better than the standard BM25 algorithm.

The query sentences were preprocessed using the NLTK (Natural Language Toolkit) package [34]. The sentences were first tokenized into words. Greek alphabets (α, β …) were converted into English words (alpha, beta, etc.). Stop words and punctuations were removed. The remaining tokens were rejoined with a space and used as the final query to the databases. When searching PubMed, we added “OR” between each pair of consecutive words. This is necessary for PubMed to return results. Sentences with less than 5 tokens or more than 50 tokens after preprocessing (tokenization, conversion of Greek alphabets, removal of stop words and punctuations) were excluded. For each sentence query, we obtained the top 1000 relevant articles among all PubMed articles using MBM, PubMed TF-IDF (PMT) algorithm (the old PubMed algorithm before Best Match) or PubMed BestMatch (PMB) algorithm. Query sentences, whose corresponding cited articles have less than 50 words in their titles and abstracts, were excluded.

To develop the training and validation datasets, the query sentences were paired with their corresponding cited articles and these pairs served as positive cases. In addition, two negative cases were constructed for each citing sentence by pairing the citing sentence with two articles not cited by the sentence: one randomly selected from the top 1,000 search results of the citing sentence and another randomly selected from all other articles. In total, there are 854,101 sentences with 936,591 citations paired as positive cases and 1,870,387 negative cases in the training data. There are 145,455 sentences paired with 148,269 citations as positive cases and 296,128 citations as negative cases in the validation dataset.

Three test datasets were developed to evaluate different methods. The test datasets were constructed by querying the citing sentences against all PubMed abstracts using MBM, PMT, or PMB. Biopython package [35] was used to query the PubMed database. We first randomly selected 90,757 sentences whose cited articles are ranked in the top 1000 of search results by MBM. This test dataset is referred to as D1. More details on dataset D1 are given in Table S1 (Supplementary Materials).

The second test dataset (D2) includes cases for which the articles cited by the query sentences were all ranked in top 1000 by both MBM and PMT. The PubMed database implemented two relevance scoring algorithms: the TF-IDF algorithm has been used since 2013 and the “Best Match” algorithm, which is a L2R (learning to rank) machine learning algorithm, was deployed in 2017. Querying through the NCBI Entrez API gives results ranked by the TF-IDF algorithm while querying through the PubMed web portal gives results ranked by the “Best Match” algorithm. We used the Biopython package [35] to query the PubMed database through the Entrez API to get search results ranked by the TF-IDF algorithm, and wrote a Python script to query the PubMed database through its web API to get search results ranked by the “Best Match” algorithm. Among the 90,757 citing sentences, 57,123 sentences had their corresponding cited articles ranked in the top 1,000 search results by both MBM and PMT. Table S2 (Supplementary Materials) provides more details on dataset D2.

Querying the PubMed database through its web API was much less efficient than through the Entrez API. We managed to get search results for 9,916 citing sentences for which the cited articles were ranked in the top 1,000 search results by MBM, PMT, and PMB. This dataset is referred to as dataset D3 (Table S3 in Supplementary Materials).

To investigate the performance on sentence queries of Google Scholar, we randomly selected 100 query sentences from D3. The sentences were used as queries to manually search on Google Scholar. In all the test datasets, the cited article of a query sentence is considered as the true relevant article for that sentence, and we check whether a search method ranked it in the top 1, 20, or 100 among all the 1,000 search results.

### The deep learning model

After experimenting with different deep learning models including BERT (Bidirectional Encoder Representations from Transformers) [36], we found that the decomposable attention model [37] achieved the best tradeoff between accuracy and speed. The decomposable attention model is a simple neural architecture proposed by Parikh et al. for natural language inference. The core of the architecture consists of three steps: Attend, Compare, and Aggregate. To accommodate the characteristics of the search engine, a modified decomposable attention model was implemented in this work. Architecture of the modified decomposable attention model is shown in Figure 2.

**Figure 2.**
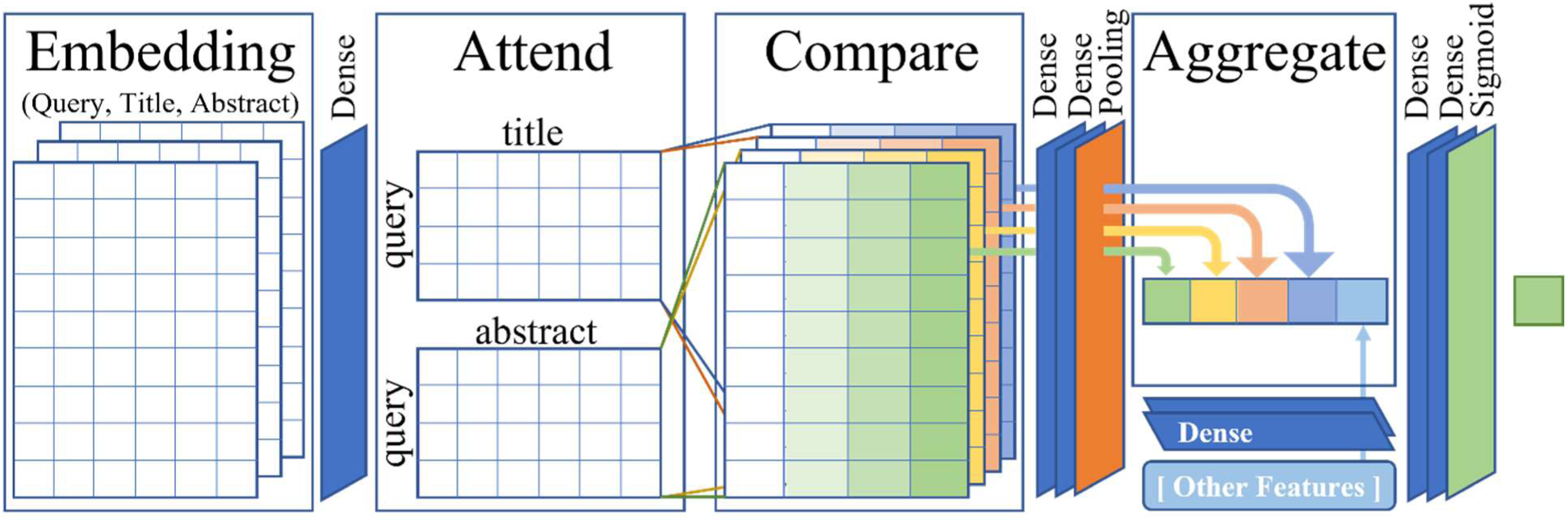
The Decomposable Attention Model.

At the training stage, the inputs are the quadruplets 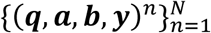, where ***q*** is the query sentence, ***a*** and ***b*** are the title and abstract of the cited article, and ***y*** is the label. At the evaluation stage, each query sentence was paired with each of the top returned articles. The triplet (***q, a, b***) was used as the input to the model that predicted the probability of relevance ranking for the returned article.

#### Embedding

Word embeddings with dimension *d* were used as representation of inputs. Let 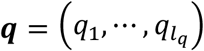 be the query sentence, 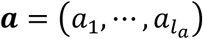 and 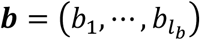 be the title and abstract of the cited article or returned article, respectively. *l*_*q*_, *l*_*a*_, *l*_*b*_ are the lengths of the query sentence, the title, and the abstract of the article, respectively. *q*_*i*_, *a*_*i*_, *b*_*i*_ ∈ ℝ^*d*^ are the *i*^th^ word of the query sentence, the title, and the abstract of the article. The inputs are passed through a dense layer *F* with an exponential linear unit (ELU) [38] activation function and dropout. The resultant is then fed to subsequent steps of the model.

#### Attend

At the Attend step, we first compute the unnormalized attention weights 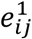 for each element of the query sentence and the title, and for each element of the query sentence and the abstract.

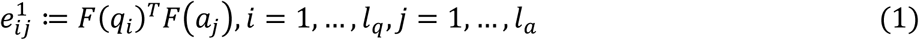

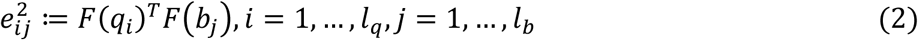

Then the normalized attention representations, 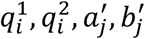, can be calculated as follows:

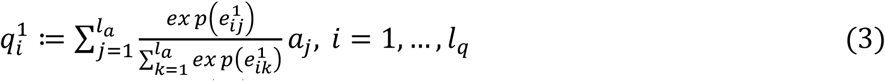

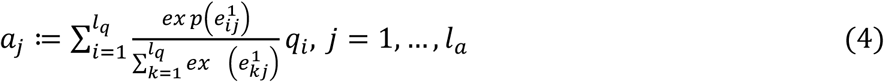

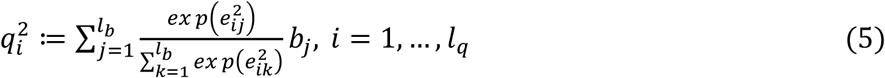

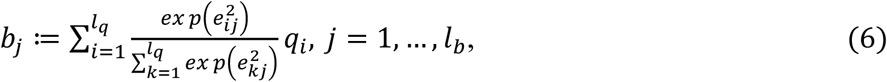

where 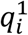 and 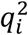 are the normalized attention representations of the query sentence attending to the title and abstract of the article, and 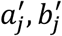 are the normalized attention representations of the title and abstract of the article attending to the query sentence.

#### Compare

At the Compare step, we concatenate the corresponding input representation, the normalized attention representation, and the difference between the input representation and the normalized attention representation for the query sentence, the title and the abstract. Then a fully connected layer *G*, with ELU activation function, dropout and max pooling was applied on top of the concatenation as follows:

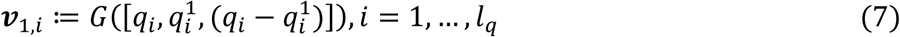

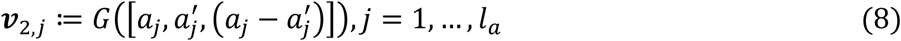

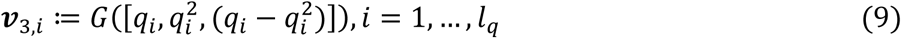

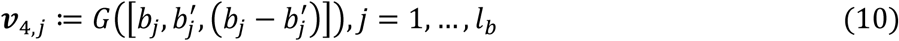

#### Aggregate

At the step of Aggregate, we first aggregate elementwise the representations of the query sentence, the title and the abstract through summation as follows:

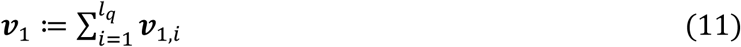

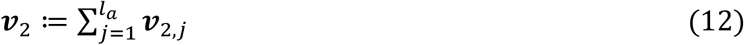

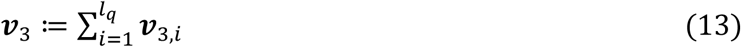

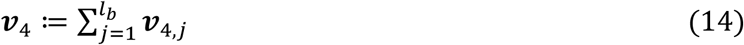

Additional features are concatenated and passed through two fully connected layers with an ELU activation function, dropout, and Batch Normalization [39] for an aggregated representation (function *L*() in equation 15).

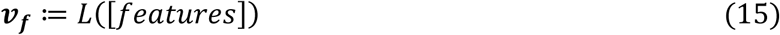

All aggregated representations are then concatenated and passed through two fully connected layers with ELU activation function, a dropout, a Batch Normalization, and finally a Sigmoid layer (function *H*() in equation 16).

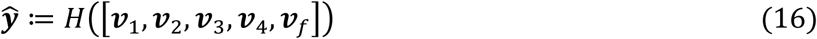

### Model training

The modified decomposable attention model was trained and validated on the training and validation datasets using the following settings: word embedding dimensions: 300; numbers of hidden layers: [500, 300, 32, 16]; dropout rate: 0.2; optimizer: Adam [40]; loss function: binary cross-entropy; learning rate: 0.01; batch size: 20; max training epochs: 20. We also used a 300-dimensional word embedding trained on more than 17 million PubMed articles using fastText [41]. The maximum length of query sentences and article titles was set to be 100 words, while the maximum length of article abstracts was set to be 1,000 words. Early stopping was used to determine the best epoch by monitoring the loss on the validation dataset. The programs used in this study were implemented in Python. We used Keras [42] together with TensorFlow[43] to implement deep learning models.

## Results

We first evaluated DeepSenSe using test dataset D1. In D1, there are 90,757 sentences, and each sentence has 1000 candidate relevant articles including the article the sentence actually cited. For each query sentence, we check whether the algorithm can rank the corresponding cited article in the top *k* articles among the 1000 candidate articles, where *k* = 1, 10, and 100. To compare ranking algorithms, we compare the numbers of times the ranking algorithms can rank the cited articles in top *k* (*n*_*top-k*_) for all 90,757 cases. For MBM the *n*_*top-k*_ values are 17,898, 44,957, and 62,609 for *k* = 1, 10, and 100, respectively. As a comparison, DeepSenSe was able to improve these values to 23,649, 61,132, and 79,830, respectively. The relative increases are 32%, 36% and 28%, respectively (Figure 3A).

**Figure 3:**
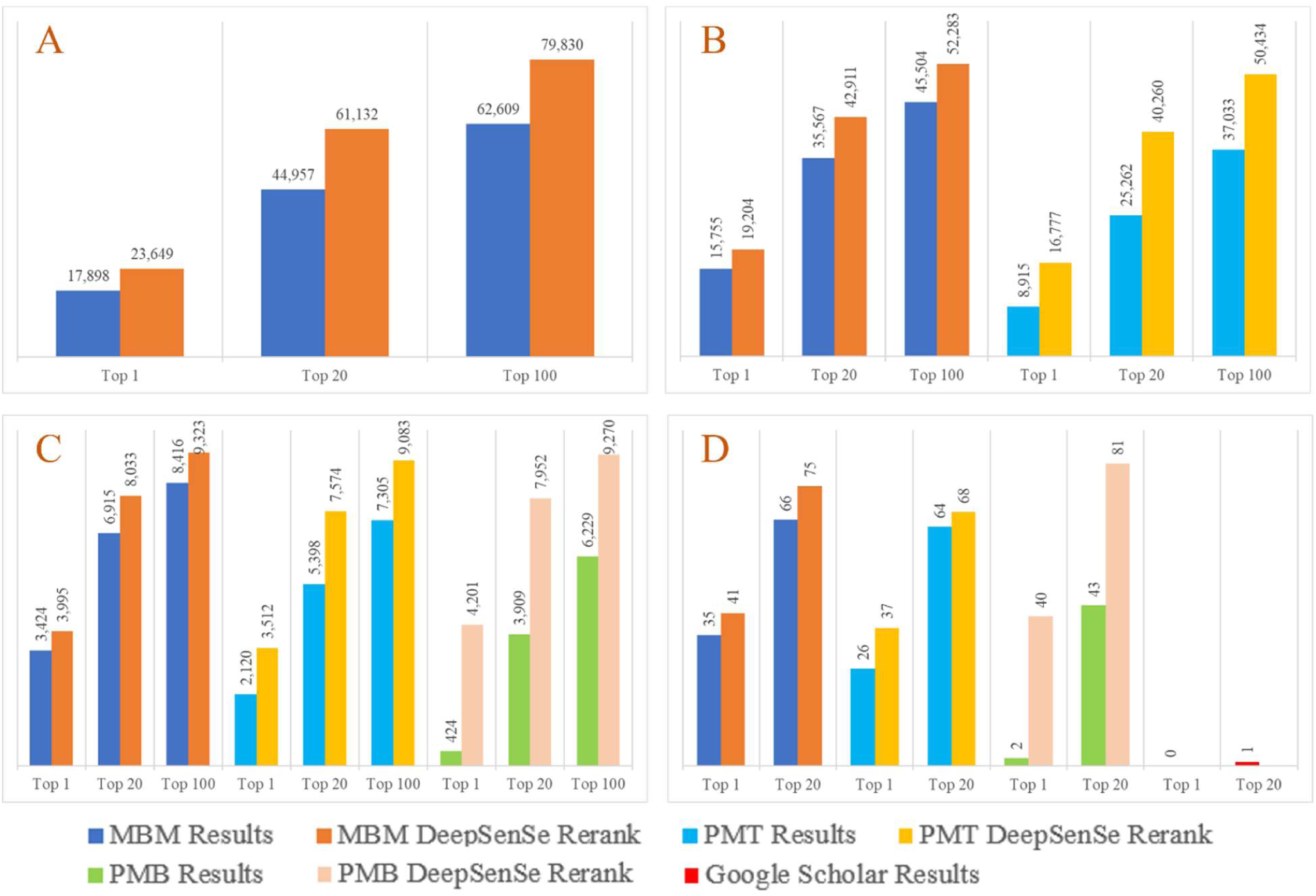
Performance of DeepSenSe, MBM, PMT, PMB, and Google Scholar. A: Performance of DeepSenSe and MBM on D1; B: Performance of DeepSenSe, MBM and PMT on D2; C: Performance of DeepSenSe, MBM, PMT and PMB on D3; D: Performance of DeepSenSe, MBM, PMT, PMB and Google Scholar on 100 sentences randomly selected from D3. Top1: the relevant article is ranked as top 1 of the search result; top20: the relevant article is ranked within the top 20 of the search result; and top100: the relevant article is ranked within the top 100 of the search result. MBM: modified BM25; PMT: PubMed TFIDF; PMB: PubMed Best Match.

We compared DeepSenSe with PubMed TF-IDF (PMT) algorithm using D2 with 57,123 query sentences, in which the articles cited by the query sentences were all ranked in top 1000 by both MBM and PMT. The top 1000 articles from MBM and PubMed are different. So, we let DeepSenSe to re-rank both sets of 1000 articles for each query sentence. Compared to MBM, the DeepSenSe model was able to improve the *n*_*top-k*_ values from 15,755 to 19,204, 35,567 to 42,911, and 45,504 to 52,283 for *k* = 1, 10, and 100, respectively. The relative increases are 22% (k=1), 21% (k=10), and 15% (k=100). Compared to PMT, DeepSenSe improved the values from 8,915 to 16,777 (k=1), 25,262 to 40,260 (k=10), and 37,033 to 50,434 (k=100). The relative increases are 88%, 59%, and 36%, respectively (Figure 3B).

We then compared DeepSenSe with PMT and PubMed BestMatch (PMB) algorithms. We used D3 with 9,916 sentences, where the articles cited by the query sentences were all ranked in top 1000 by MBM, PMT, and PMB. Again, DeepSenSe was able to improve the ranks substantially for all three algorithms for this dataset (Figure 3C).

Finally, we compared DeepSenSe with PMT, PMB, and Google Scholar using a very small test dataset with only 100 sentences, since Google does not allow automatic querying of their system. These 100 sentences were randomly selected from dataset D3. The comparison results are shown in Figure 3D. In the Google Scholar search, we limited the domain to PubMed database to be consistent with other searches. The performance of Google Scholar for this dataset is much worse than the other search algorithms. Again, DeepSenSe performed the best among all the search algorithms.

In Table S4 (Supplementary Materials), we show some examples, which DeepSenSe and MBM ranked very differently. A clear trend we have observed is that DeepSenSe matches meanings better than MBM, which, as a BM25 based method, matches exact keywords better.

## Conclusions and Discussions

Document retrieval using full sentences as queries can help users find relevant documents more effectively. It is also very useful for building question-answering systems, identifying relevant citations for scientific manuscripts, and comparing new findings with previous knowledge. In this study, we developed a deep learning model, called DeepSenSe, trained using a large volume of labeled data obtained from the citation data of PMC full-text articles. Tested on large test datasets, DeepSenSe was able to substantially improve the rankings of existing methods including a modified BM25 (MBM) and PubMed’s ranking algorithms. The combination of MBM and DeepSenSe gave the best performance overall.

We tried several different deep learning architectures and the decomposable attention model had the best tradeoff between accuracy and speed (in terms of both training and prediction). The BERT model had better performance, but is much slower in training and prediction. With more powerful hardware, it is possible that more sophisticated models can be employed to achieve even better overall performance in the future.

User behavior analysis on PubMed showed that most queries are short and over 80% of all queries had no more than four tokens [44-46]. We hypothesized that this may be partially due to the current search engines not performing well for long queries, so that users do not tend to use them as much. This user behavior could change if they find long queries can give them better results.

In this study, we focused more on whether a relevant article will rank in the top 1, 20, and 100, instead of its absolute rank. Since over 80% of users only clicked on the results from the first page [44], it is crucial to show the most relevant articles on the first page. If a page shows 20 results then being ranked in the top 20 means the article is on the first page, and it has a much higher chance to be found by a user.

We used ELU instead of ReLU because the effect of words with opposite meanings was also significant in our situation. If we do not use ELU, the model predicts very high relevance scores for almost all search results so that we cannot find out the cited articles.

It is worth mentioning that MBM, PubMed, and Google Scholar were not optimized for sentence queries. So, the comparison in this study simply showed that DeepSenSe can improve MBM and PubMed ranking results substantially for sentence queries using citation data as tests, instead of a demonstration that DeepSenSe is better than these methods for general queries. Applying the same concept to developing a search engine for general queries will be the subject of future studies.

In addition to providing more relevant search results for sentence queries for search engines, DeepSenSe is also ideally suited to scan full-text documents to identify relevant citations for the sentences in the documents. For example, they can help authors add citations to the articles they are writing or help institutions/agencies with document review by assessing the quality and completeness of the citations in the documents. We are currently developing such a system, which will be released in late 2021.

## Acknowledgement

JZ is supported partially by a grant from National Institute of General Medical Science of National Institutes of Health, grant No. R01GM126558. The funder had no role in the study design, data collection and analysis, decision to publish, or preparation of the manuscript.

## Notes

### Competing Interest Statement

The authors have declared no competing interest.

