## Supplementary Material for "Developing a More Accurate Biomedical Literature Retrieval Method using Deep Learning and Citations in PubMed Central Full-text Articles"

DeepSenSe: A Deep Learning Method for Searching Biomedical Literature using Sentences as Queries

Table S1. Summary of the Large Test Dataset, D1.

| | | MBM |
|---|---|---|
| Sentences | | 90,757 |
| Ranks of Cited Articles in Query Returns | Top 1,000 | 90,757 |
| | Top 100 | 62,609 |
| | Top 20 | 44,957 |
| | Top 1 | 17,898 |

Table S2. Summary of the Medium Test Dataset, D2.

| | | MBM | PMT |
|---|---|---|---|
| Sentences | | 57,123 | |
| Ranks of Cited Articles in Query Returns | Top 1,000 | 57,123 | |
| | Top 100 | 45,504 | 37,033 |
| | Top 20 | 35,567 | 25,262 |
| | Top 1 | 15,755 | 8,915 |

Table S3. Summary of the Small Test Dataset, D3.

| | | MBM | PMT | PMB |
|---|---|---|---|---|
| Sentences | | 9,916 | | |
| Ranks of Cited Articles in Query Returns | Top 1,000 | 9,916 | | |
| | Top 100 | 8,416 | 7,305 | 6,229 |
| | Top 20 | 6,915 | 5,398 | 3,909 |
| | Top 1 | 3,424 | 2,120 | 424 |

### Example Analysis

For search engine comparison, MBM is much better than PMT and PMB. The re-rank from the deep learning model improved the accuracy significantly for all search engines. Table S4 showed some examples for which DeepSenSe had very different rankings from MBM.

Table S4. Examples analysis for search engine and deep learning model results.

| Category | Search Rank | Model Rank | SEN PMID | Sentence (SEN) | ABS PMID | Cited Article's Title (ABS) |
|---|---|---|---|---|---|---|

| | | | | | | |
|---|---|---|---|---|---|---|
| A-1 | not found | 201 | 29739739 | Use of K-SRS assessment research (n=13) was done to study system effectiveness, feasibility, accuracy, or reliability. | 25884838 | Instrumenting Gait Assessment Using the Kinect in People Living With Stroke: Reliability and Association With Balance Tests |
| A-2 | not found | 711 | 28381227 | All but one study (examining elderly) reported overweight/obesity to be associated with lower levels of individual education | 24133894 | Association Between Obesity and Symptoms of Depression of Adults in Puerto Rico |
| B-1 | not found | 1 | 27135338 | Put and Spd are essential for life, as Arabidopsis mutants defective in their biosynthetic pathways are embryo-lethal, whereas Spm and T-Spm have been specifically linked to stress responses and development, respectively. | 18594857 | Polyamines: Essential Factors for Growth and Survival |
| B-2 | not found | 10 | 27382330 | These are the main mechanisms leading to a prolongation of action of aminosteroid NMBAs. | 1677546 | Distribution, Elimination, and Action of Vecuronium in the Elderly |
| C | 1 | 290 | 28321231 | Proline accumulation is considered to be a plant adaptive response to high salinity and drought stresses. | 21400017 | Drought-induced Proline Accumulation Is Uninvolved With Increased Nitric Oxide, Which Alleviates Drought Stress by Decreasing Transpiration in Rice |
| D-1 | 891 | 1 | 19590675 | The prevalence of African women who habitually use Hg-containing cosmetic products is not known; however, surveys in certain populations suggest that the use of these cosmetic products is quite variable, ranging from 10% in a study of Senegalese women to 47% in a population of male and female Nigerian traders. | 12081345 | An epidemiological survey of the use of cosmetic skin lightening cosmetics among traders in Lagos, Nigeria. |
| D-2 | 296 | 4 | 29142642 | As having a hydrophobic methacrylate terminal end and a hydrophilic phosphate terminal end, copolymerizing resin monomers and chemically binds to oxides, respectively, MDP has a bifunctional adhesive monomer that can bind to zirconia or metal. | 16193486 | Bonding of Dual-Cured Resin Cement to Zirconia Ceramic Using Phosphate Acid Ester Monomer and Zirconate Coupler |
| D-3 | 288 | 4 | 19847088 | Similarly all the three patients in the study by Holmang et al with focal LELC died of the disease (9-68 months), compared to none of the six patients with pure or predominant LELC (13 months to 18 years) | 9474147 | Bladder Carcinoma With Lymphoepithelioma-Like Differentiation: A Report of 9 Cases |
| D-4 | 680 | 14 | 27435901 | Hydrophobic interactions are commonly described, but since cognate domains are selective to their partner PCP(s), further interactions, such as variable charge distribution,20 must also play important roles in recognition. | 25050442 | The Crystal Structure of BlmI as a Model for Nonribosomal Peptide Synthetase Peptidyl Carrier Proteins |

| | | | | | | |
|---|---|---|---|---|---|---|
| D-5 | 665 | 6 | 23698162 | There is general agreement that increasing dietary SFA intake, especially in overweight or obese individuals, is associated with raised inflammatory markers, predominately by activating the toll-like receptor 4 (TLR4) pathway. | 22133051 | Dietary Factors and Low-Grade Inflammation in Relation to Overweight and Obesity |
| D-6 | 192 | 4 | 29593498 | The subventricular zone (SVZ) is a prime neuropoietic niche of the brain responsible for the postnatal neurogenesis in the telencephalon. | 10380923 | Subventricular Zone Astrocytes Are Neural Stem Cells in the Adult Mammalian Brain |
| D-7 | 158 | 3 | 29376513 | Especially educational measures, community based measures as well as legislative measures were set up to reduce injuries in the young population. | 9346041 | Can We Prevent Accidental Injury to Adolescents? A Systematic Review of the Evidence |
| D-8 | 863 | 5 | 26035426 | The size of mtDNA nucleoids is debated in the literature (although recent evidence from high-resolution microscopy suggests that nucleoid size is generally <2, consonant with recent evidence that individual nucleoids may be homoplasmic); our model allows for inheritance of homoplasmic or heteroplasmic nucleoids of arbitrary characteristic size c, thus allowing for a range of sub-organellar mtDNA structure. | 23721879 | mtDNA makes a U-turn for the mitochondrial nucleoid. |
| D-9 | 961 | 8 | 23176133 | A modified Borg scale of perceived exertion will be used to regulate the intensity of resistance exercise. | 7154893 | Psychophysical bases of perceived exertion. |

Note: Because abstracts are too long, we cannot show them here. Those abstracts can be found by searching ABS PMID on PubMed.

Table S4 is an analysis of the MBM search engine and the deep learning model results. In category A, both the MBM search engine and the model have bad results because SENs do not contain terminologies. In category B, the MBM search engine cannot find common words. However, our model can catch the similarity of related words, such as Spd, Spm, and Polyamines. In category C, it was hard to distinguish the cited article from others because those articles contained many words in SEN like proline accumulation, plant, and stress. The PMID of the rank 1 article in our model was 18379856. In category D-1, the MBM search engine did not have a good result because SEN used African and Nigerian, but ABS used Nigeria. Our model was able to know they were related. The PMID of the rank 1 article in the search result was 10805050, which contained African and Nigerian. In D-3, Sentence only mentioned LELC, which is an abbreviation of Lymphoepithelioma-like carcinoma.
